# Optimising experimental designs for model selection of ion channel drug binding mechanisms

**DOI:** 10.1101/2024.08.20.608856

**Authors:** Frankie Patten-Elliott, Chon Lok Lei, Simon P. Preston, Richard D. Wilkinson, Gary R. Mirams

## Abstract

The rapid delayed rectifier current carried by the human Ether-à-go-go-Related Gene (hERG) channel is susceptible to drug-induced reduction which can lead to an increased risk of cardiac arrhythmia. Establishing the mechanism by which a specific drug compound binds to hERG can help to reduce uncertainty when quantifying pro-arrhythmic risk. In this study, we introduce a methodology for optimising experimental voltage protocols to produce data that enable different proposed models for the drug-binding mechanism to be distinguished. We demonstrate the performance of this methodology via a synthetic data study. If the underlying model of hERG current is known exactly, then the optimised protocols generated show noticeable improvements in our ability to select the true model when compared to a simple protocol used in previous studies. However, if the model is not known exactly, and we assume a discrepancy between the data-generating hERG model and the hERG model used in fitting the models, then the optimised protocols become less effective in determining the ‘true’ binding dynamics. While the introduced methodology shows promise, we must be careful to ensure that, if applied in a real data study, we have a well-calibrated model of hERG current gating.

## 1 Introduction

Ion channels are proteins in the cell membrane that form pores through which ions can flow in and out of the cell. The resulting ion currents play an important role in several biological functions including coordinating the contraction of muscle cells. A healthy heart relies on regular, coordinated contractions of cardiomyocytes (heart muscle cells) to pump blood from the heart around the body [1]. The K_v_11.1 ion channel encoded by the human Ether-à-go-go-Related Gene (hERG) is responsible for conducting the rapid delayed rectifier potassium current, *I*_Kr_, and plays a crucial role in cardiomyocytes recovering from excitation [2]. However, the hERG channel is susceptible to block by pharmaceutical small molecules (referred to here as ‘compounds’ throughout); this can lead to a reduction in *I*_Kr_, lengthening the cardiac action potential, and, in some cases, increasing the risk of cardiac arrhythmia [3, 4, 5].

Markov-style computational models of ion channels define transition rates between several channel states (e.g. open, inactive, closed) and can be used to simulate channel current in response to a membrane potential. To model the interactions of drug compounds with an ion channel, additional states and rates can be introduced to an existing ion channel model to simulate various binding mechanisms [6]. When it comes to the hERG channel, it has been observed that the binding mechanism can be compound-specific [7, 8, 9]. One such example of this is the propensity of a compound to become ‘trapped’ (unable to unbind) when the channel closes. Some compounds, such as bepridil and dofetilide, are known to become trapped inside the central hERG cavity, remaining bound, while others, such as cisapride and verapamil, unbind when the channel closes [10, 11, 12]. It has also been theorised that some drugs bind preferentially to certain channel states over others, or in the extreme case, bind to a particular state only [13, 14, 15, 16, 17]. These compound-specific binding mechanisms suggest that a one-size-fits-all approach to modelling hERG-drug interactions is perhaps limiting [6]. To accurately model how a certain compound binds to the hERG channel it is, therefore, important to determine the specific mechanisms at play.

The transmembrane current of a cell can be measured in response to the membrane potential via a voltage-clamp experiment where a piecewise function defining the transmembrane voltage, *V*, dependent on time, *t*, is applied. We will refer to this function, *V* (*t*), as a *voltage protocol*. In a recent study, Lei *et al*. [18] considered fitting a set of 15 pharmacological models representing possible drug-binding mechanisms (trapping/non-trapping, state binding preference, etc.) to previously collected voltage-clamp data under a relatively simple protocol. After fitting the set of models, it was suggested that more information-rich experiments may be needed to distinguish between model outputs and to assist in determining compound-specific binding mechanisms. Previous studies on drug-binding dynamics have considered ‘manual’ experimental design techniques to increase the information extracted from voltage-clamp experiments [19, 20, 21, 22, 23]. This often involves some degree of expert knowledge to design protocols that are expected to emphasise particular compound-specific behaviours. In this work, we instead consider ‘automated’ Optimal Experimental Design (OED) techniques.

OED methods consider how the design of a data-collecting experiment can be optimised with respect to some statistical criterion, effectively maximising the information provided by the experiment (subject to constraints). These methods have been used recently in the field of cardiac modelling with some success [24]. In this paper, we consider OED methods to design voltage protocols that can be used in voltage-clamp experiments to better distinguish between different models of drug-binding mechanism. We detail a synthetic data methodology for generating an optimised protocol and fitting models to data collected under this protocol. Our results demonstrate how the optimised designs can assist in establishing the true binding dynamics at play across a range of simulated drug compounds exhibiting differing dynamics. However, we find that introducing a *discrepancy* between the hidden ‘true’ data-generating model and the proposed model we work with to fit the data reduces the effectiveness of this method, and suggests that further work may be needed to account for these inevitable model discrepancies when working with real data.

## 2 Methods

### 2.1 Mathematical models

#### hERG physiological models

In this paper, we consider two physiological models of hERG; one simple four-state model used throughout, and a slightly more complex five-state model exhibiting differing behaviour that is used to introduce hERG model discrepancy. These models describe the voltage-dependent gating behaviour of *I*_Kr_ at physiological temperature when no drug compound is present. Figure 1 shows Markov diagrams of the two hERG models we will be considering. Figure 1a shows the four-state model, with transition rates *k*_1_ to *k*_4_, which is equivalent to physiological model B in Lei *et al*. [18]. The four states in this model are IC (Inactive Closed), C (Closed), I (Inactive), and O (Open), where each variable represents the proportion of channels in that state. Figure 1b shows the five-state model from Lu *et al*. [25], with transition rates *a*_*a*0_, *b*_*a*0_, *k*_*f*_, *k*_*b*_, *a*_*a*1_, *b*_*a*1_, *a*_*ci*_, *b*_*ci*_, *a*_*i*_, and *b*_*i*_. This model has three closed states (C1, C2, C3), and no states that are both inactive and closed. All transition rates in both models (apart from *k*_*f*_ and *k*_*b*_) are dependent on transmembrane voltage, *V* ; they are defined by the general equation *p*_*i*_exp(*p*_*j*_*V*) where *p*_*i*_ and *p*_*j*_ are physiological model parameters taken from the literature. In both models, the rate of change of each state over time, *t*, is defined by a differential equation. For example, for the model in Figure 1a, the rate of change of open proportion is defined by

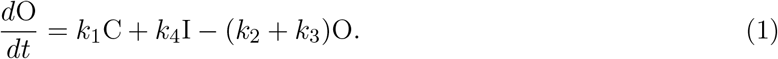

**Figure 1:**
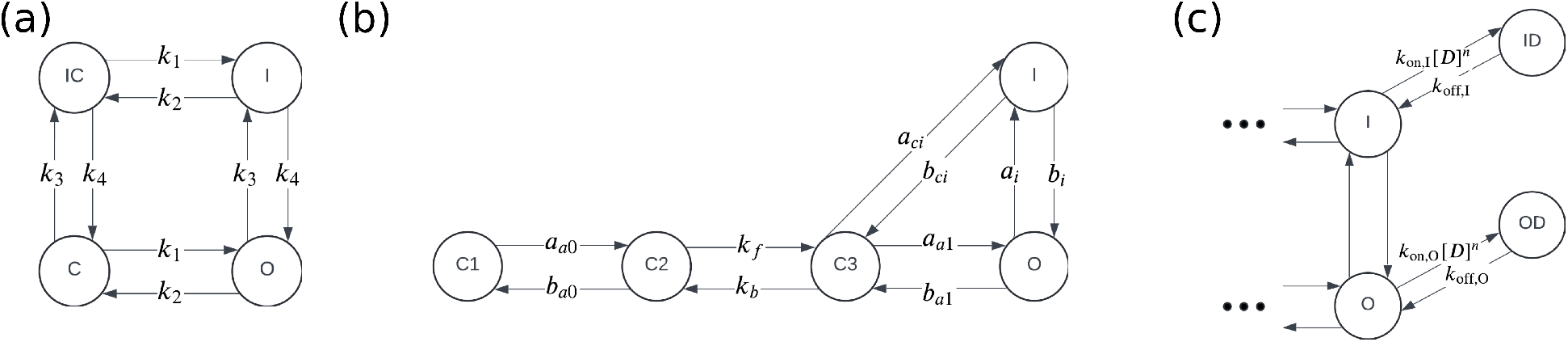
Markov diagrams of physiological hERG models (a, b) and a pharmacological binding model (c). The model in (a) is a four-state symmetric hERG model as used by Lei *et al*. [18]. The four states are IC (Inactive Closed), C (Closed), I (Inactive), and O (Open). The model in (b) is a five-state model equivalent to that derived by Lu *et al*. [25]. This model has three closed states (C1, C2, and C3) as well as O (Open) and I (Inactive) states. The pharmacological drug-binding in (c) is Model 7 from Lei *et al*. [18]. The I and O states represent the inactive and open states in the underlying physiological hERG models (this model could be attached to the right-hand side of either physiological model shown in (a) or (b)). The binding model additionally has two drug-bound states ID (Inactive Drug-bound) and OD (Open Drug-bound), in which no current flows.

The measured current, *I*_Kr_, is then calculated for both models via the following equation

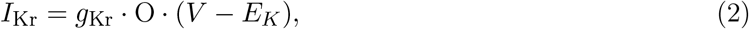

where *g*_Kr_ is conductance, O is the open state proportion, and *E*_*K*_ is the Nernst potential (membrane potential at which there is no net flux).

#### Pharmacological binding models for hERG

We can extend the Markov models of hERG described in the previous section to model drug-binding dynamics in hERG. We will consider a set of 15 models that characterise different proposed candidate mechanisms for drug-binding as illustrated in Figure 2 in Lei *et al*. [18] and included in the supplementary material in Figure S1. Figure 1c shows a Markov diagram of drug-binding Model 7 as an example of one of these 15 models. In this example model, we have two additional states: ID (Inactive Drug-bound) and OD (Open Drug-bound). Binding rates are described by the parameters *k*_on,I_, *k*_off,I_, *k*_on,O_, and *k*_off,O_ and the rate of on-binding is dependent on the drug concentration [*D*] and the Hill coefficient *n*.

**Figure 2:**
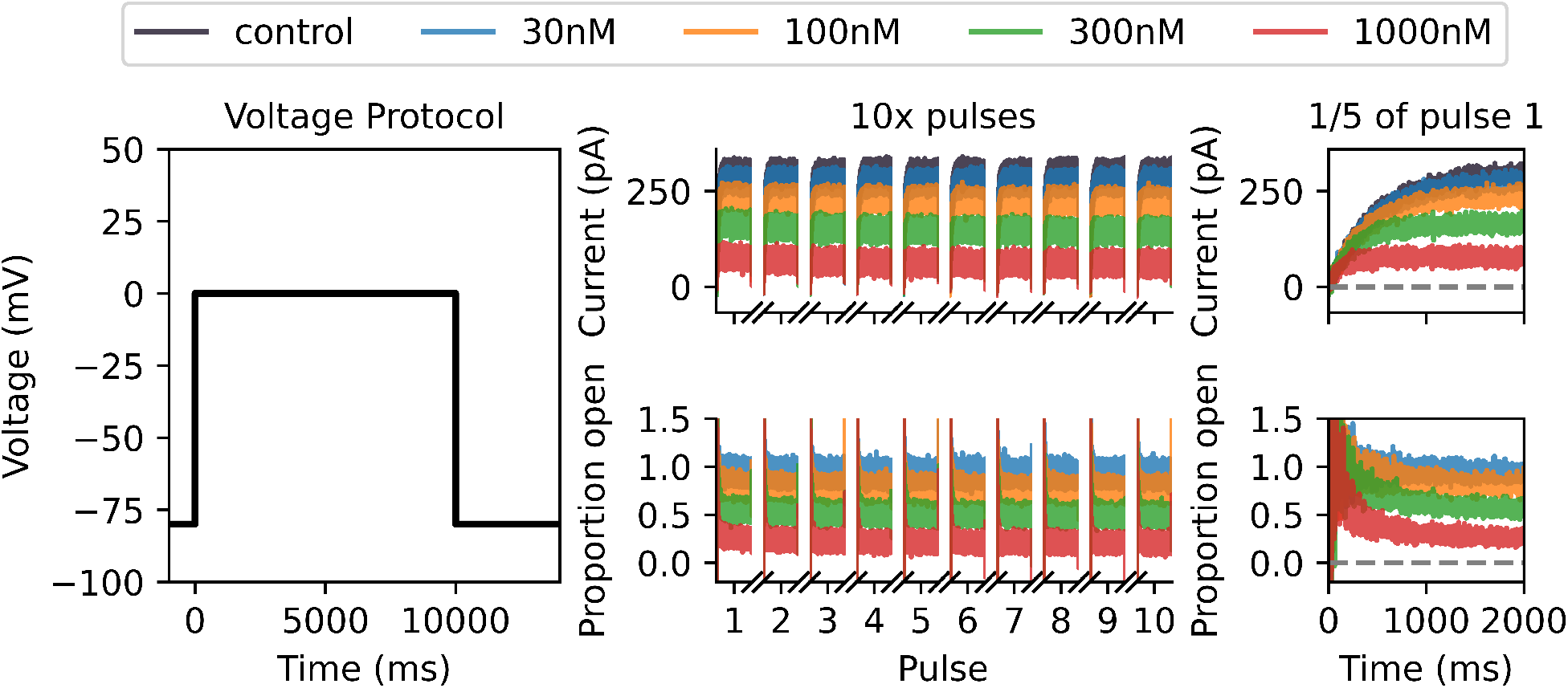
Synthetic verapamil data under drug-binding Model 7 generated under a Milnes protocol with the Lei 37°C hERG model. At the left, we plot a single sweep of the Milnes protocol. In the top middle, we plot the control (in black) and drug currents (blue, orange, green, and red corresponding to four different drug concentrations) for 10 sweeps of the Milnes protocol. We only plot the currents that occur during the 10s pulse at 0 mV for each sweep. In the bottom middle, we plot the corresponding proportion open for each drug concentration which is calculated by dividing each drug sweep by the control current. On the right, we include a zoomed-in look at the first 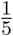 (2 s) of the first pulse for both currents and proportion open. This illustrates the currents starting at approximately 0 pA, and the open proportion starting at approximately 1. The noise in the initial low currents contributes to the increased noise at the beginning of the open proportion sweeps. This motivates using a noise model for the ratio of two normally distributed variables when fitting drug-binding models to this data.

### 2.2 Initial synthetic data

To measure the effect of drug block on the hERG channel, we can collect two sets of voltage-clamp time-series data; the control current, *y*(*t*), before the drug compound is introduced, and the current in the presence of the drug compound, *x*_*c*_(*t*) (often at several different concentrations *c* ∈ *C*). We begin by generating some synthetic ion channel drug-binding data in a form resembling what we would expect to collect in a voltage-clamp experiment under a simple modified Milnes voltage protocol as used in the Comprehensive in-vitro Proarrhythmia Assay (CiPA) initiative [19, 26]. With this synthetic data, we will fit the set of 15 drug-binding models from supplementary Figure S1, and illustrate a need for a more complex protocol to differentiate between these models.

In Figure 2 we plot the Milnes protocol, i.e. controlled voltage time-series (left), synthetic current data (top middle), and synthetic open proportion data (bottom middle). In the synthetic current plot, the control current in black, *y*(*t*), is generated under the Lei Markov model described in Figure 1 with parameters for physiological temperatures taken from Lei *et al*. [27]. We run 10 sweeps (repeats in series) of the Milnes protocol and plot the 10 s of each sweep corresponding to the 0 mV pulse in the protocol. We generate equivalent currents in the presence of four concentrations of verapamil (*x*_*c*_(*t*), *c* ∈ *C* = {30, 100, 300, 1000}) and these are plotted in blue (30 nM), orange (100 nM), green (300 nM), and red (1000 nM). Gaussian random noise is added to all sweeps with a standard deviation of 10 pA. We use a step size of 0.5 ms between data points, and we can define *T* as the set of all times for the plotted traces. The plotted proportion of channels open, *z*_*c*_(*t*), in the bottom middle of Figure 2 is calculated by dividing the drug current sweeps, *x*_*c*_(*t*), by the control sweep, *y*(*t*), and this is the normalised quantity we will use to fit the drug-binding models, as in [28, 26].

As an example for many of the figures in this paper, for our ‘true’ data-generating model of drug-binding dynamics, we have used the drug-binding Model 7 described in Lei *et al*. [18] as fitted to verapamil voltage-clamp data collected at physiological temperatures by Li *et al*. [28, 26]. Lei *et al*. [18] found that this model gave ‘plausible’ fits to the Li *et al*. data and agrees with the literature that verapamil does not tend to become trapped. This non-trapping behaviour involves having no bound closed state in the binding model; the drug can only be bound and block the channel when the channel is in an open (OD) or inactive (ID) state. Below we will go on to examine findings if any other binding model was the ‘true’ data-generating model for other drugs as well.

### 2.3 Initial model fitting

Next, we fit each of the 15 drug-binding models to the synthetic ‘proportion open’ data, *z*_*c*_(*t*). As a first pass, we assume that the underlying hERG model is known exactly (i.e. there is no hERG model discrepancy and the model parameters are known) and we wish to fit only the parameters of the drug-binding models. The data, *z*_*c*_(*t*), are generated by dividing two current traces which both have normally distributed iid noise, i.e. the data are a ratio of two normally distributed random variables. If 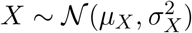 and 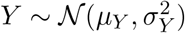 are two independent normal random variables, then the ratio *Z* = *X/Y* has probability density function (PDF) [29]

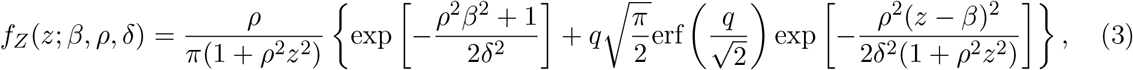

where 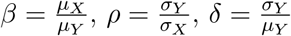, and

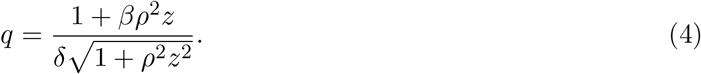

Applying this PDF to our *z*_*c*_(*t*) data case, *µ*_*X*_(*t, c, θ*) is the modelled current at time *t* in the presence of a drug compound of concentration *c* under some drug-binding parameterisation *θ*, while *µ*_*Y*_ (*t*) is the control current at time *t* and does not depend on *θ* or *c*. We will simplify things by assuming that *σ*_*X*_ and *σ*_*Y*_, the standard deviations of the measurement error on the drug and control currents respectively, are equal *σ*_*X*_ = *σ*_*Y*_ = *σ* resulting in *ρ* = 1. We can then use the PDF in equation 3 to derive a log-likelihood function for some drug-binding model parameterisation *θ* and standard deviation *σ* given a data set *z*

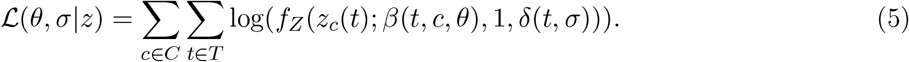

Each of our 15 drug-binding models can then be fitted to *z*_*c*_(*t*) by maximising this log-likelihood function with respect to *θ* and *σ*.

We use the covariance matrix adaptation evolution strategy (CMA-ES) optimisation algorithm [30] via the Probabilistic Inference on Noisy Time-Series (PINTS) framework [31] to perform this maximisation. We repeat the CMA-ES optimisation ten times, with each repeat starting from a different parameter initialisation sampled from wide boundaries as described in [18], and take the largest obtained log-likelihood. The choice of ten repeats is motivated by a desire to balance computation time with accuracy; on average 8 of the 10 repeats give very similar maximised likelihoods and corresponding parameter estimates. In Figure 3 we plot the fits obtained via this method for each of the 15 drug-binding models. It is difficult to visually distinguish between the quality of these fits, and, at a glance, it appears that all models fit the data relatively well. We note that the fitted parameter values obtained for Model 7 (2.19 × 10^*−*7^, 1.57 × 10^*−*5^, 5.78 × 10^*−*6^, 2.39 × 10^*−*3^, and 1.04) are very close to the true data-generating parameter values (2.19 × 10^*−*7^, 1.58 × 10^*−*5^, 5.79 × 10^*−*6^, 2.39 × 10^*−*3^, and 1.04), and the fitted value of *σ* is close to the standard deviation used to generate the synthetic data (9.40 cf. 10.0).

**Figure 3:**
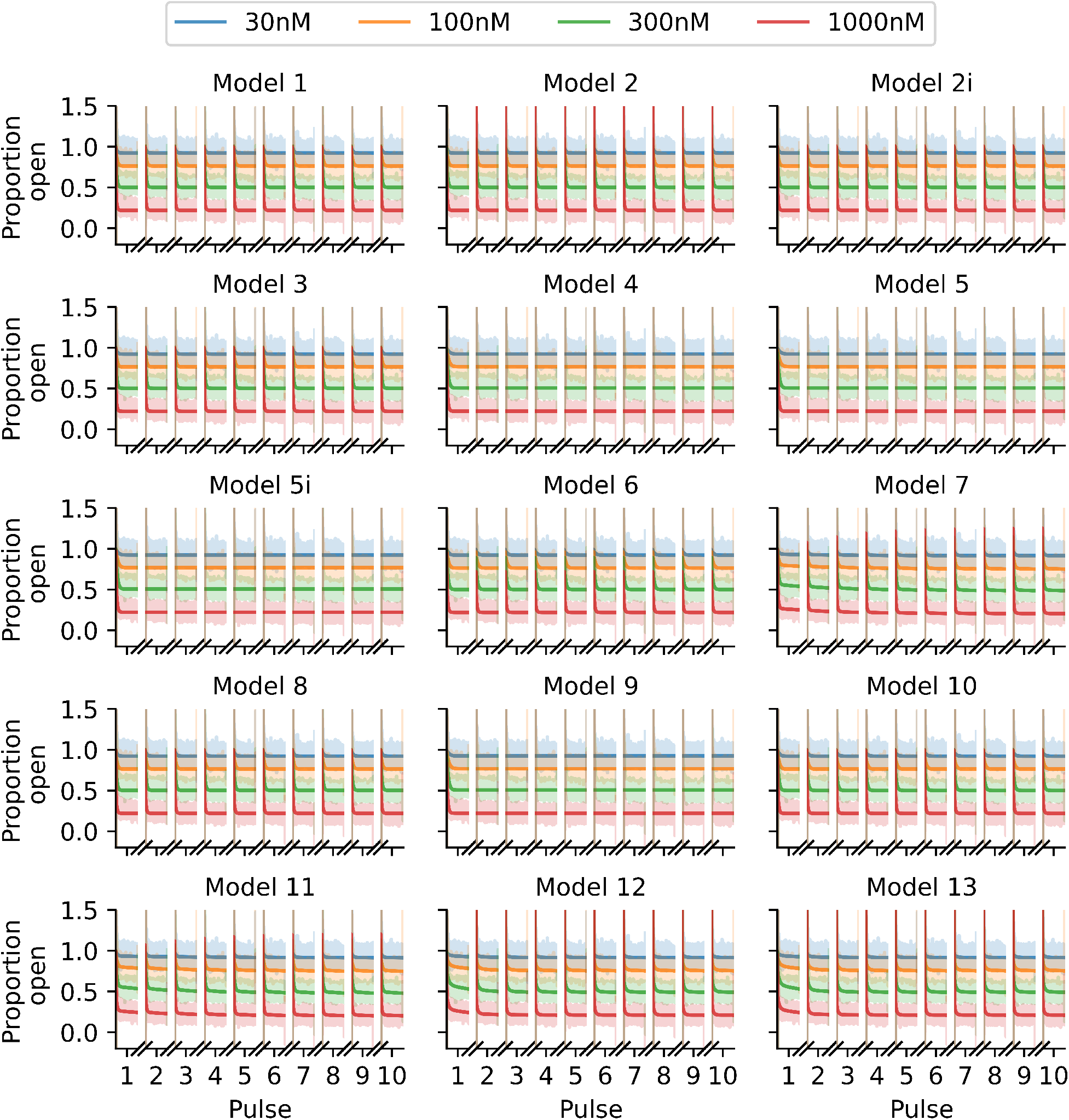
Fits of the 15 binding models to the Milnes protocol proportion-open synthetic data shown in Figure 2. Note that it is difficult to visually distinguish between the quality of the model fits for this protocol. The fitted models are shown with solid lines, and we plot the data shaded behind these fits.

In Figure 4, we plot the maximised log-likelihood values obtained when fitting each model. The true data-generating model (Model 7) has the largest maximised log-likelihood, while all other models fall within a range of 10^5^ below this value. We note that we are fitting each model to very high-resolution data (8 × 10^4^ data points), which gives rise to large log-likelihood values. A traditional model selection method such as the likelihood ratio test or the Akaike Information Criterion (AIC) may suggest, based on the differences in log-likelihoods, that the data-generating model is chosen over the other models. However, in a real data scenario, we generally have less confidence in the veracity of the model of observed hERG current due to experimental artefacts [32], so we avoid using likelihood ratio testing or AIC in this context. Ultimately, we arrive at the same conclusion drawn by Lei *et al*. [18]; this protocol is not information-rich enough to distinguish between drug-binding models, which motivates our use of OED methods.

**Figure 4:**
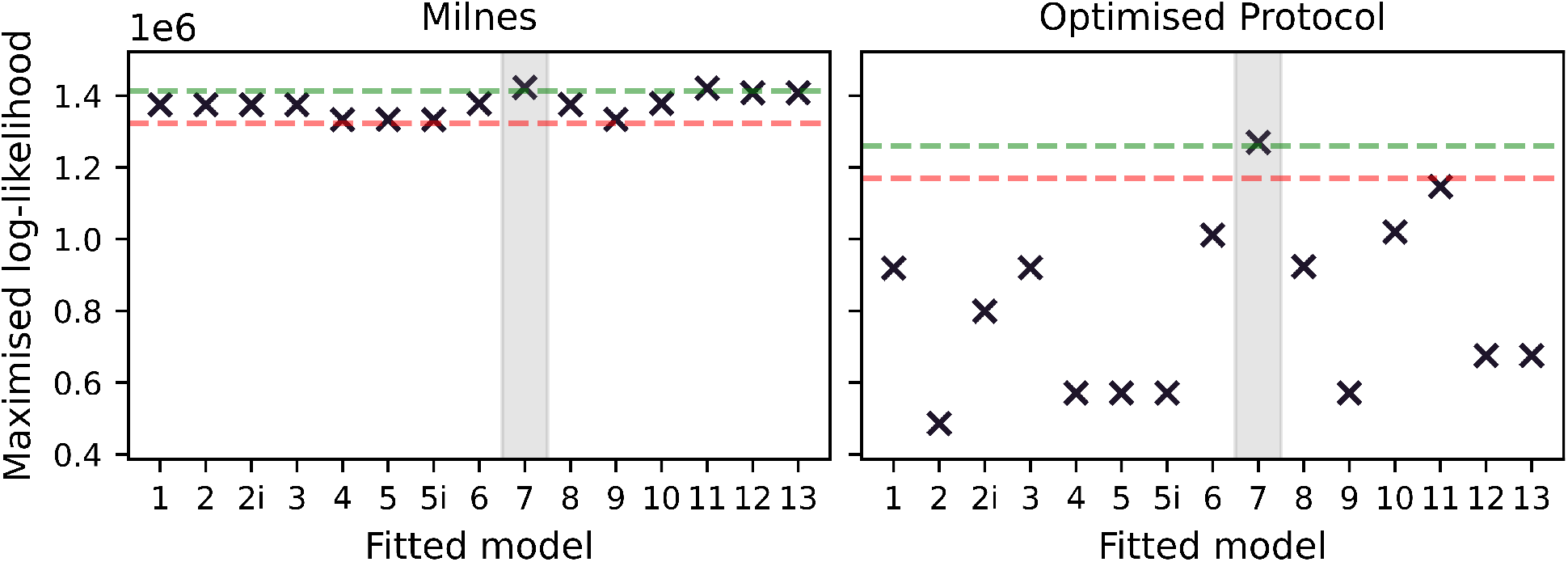
Maximised log-likelihoods for model fits to synthetic data generated under the modified Milnes protocol (left) and an optimised protocol (right). In each plot we include a green dashed line at 10^4^ below the largest maximised log-likelihood and a red dashed line at 10^5^ below the largest maximised log-likelihood. We note that these lines act simply as visual aids to emphasise the increased spread in the quality of model fits under the optimised protocol. We also shade, in grey, around the data-generating binding model.

### 2.4 Optimising the experimental design

Our next step is to develop an improved experimental design that emphasises differences between drug-binding models. We will do this by employing OED techniques. Let us assume that we have some experimental design, 𝒟, that is a function of some parameter set *ϕ*. We then need to establish some optimality criterion, *g*, that is a function of the design, and determine a *ϕ* that maximises *g*. We begin by considering *g*; what do we want to optimise? Practically speaking, we want a protocol that accentuates the differences between the models for a specific drug compound. This motivates the following optimality criterion.

#### Optimality criterion

Assume we have our fitted models from Figure 3 and these are represented by 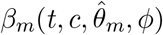 for *m* ∈ *M* where *M* is the set of 15 drug-binding models and 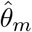 is the MLE parameter set for model *m*. Note we have included *ϕ* as an input to *β*_*m*_ here as *β*_*m*_ is dependent on the experimental design. We can then calculate the pairwise squared difference between each of the *m* model outputs, which we will label *d*_*ij*_, where

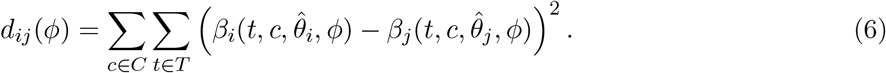

We can then propose that our optimality criterion seeks to maximise the median value of *d*_*ij*_(*ϕ*) across all *i, j* ∈ *M*, *i* ≠ *j*, which we will call *d*_*med*_(*ϕ*)

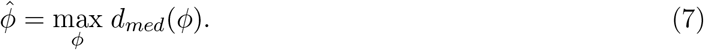

There are many choices of design criteria to optimise. A conventional option is the T-optimality criterion introduced by Atkinson & Fedorov [33], which involves maximising the *minimum* pairwise difference. However, in our case with some pairs of very similar models, initial experimentation with a T-optimality approach often resulted in the two most similar models being separated slightly, while the optimiser would ignore the pairwise differences between the other 13 models. Here we favour the *median* to ensure that the objective will seek to increase the pairwise differences between at least half of the 15 models.

#### Optimising the protocol

We now turn our attention to 𝒟(*ϕ*). With consideration of experimental practicality, we can establish a design space that we want to optimise the max-med criterion over. We begin by splitting the 10 s 0 mV pulse used in the Milnes protocol into 3 separate steps (3340 ms, 3330 ms and 3330 ms respectively) each of which can be set to a voltage in the range of −50 mV to +40 mV. To increase the design space, we include two 10 s pulses per sweep and allow both pulses to each have three different voltage steps. Further to this, we allow the times between pulses at the holding potential of −80 mV to vary; the time following the first pulse and second pulse can be set to be anywhere between 1050 ms and 21000 ms. In total, we then have eight degrees of freedom for optimising the max-med criterion; six voltage step values and two inter-pulse durations. Let us then define *ϕ* = [*V*_11_, *V*_12_, *V*_13_, *V*_21_, *V*_22_, *V*_23_, *t*_1_, *t*_2_] where *V*_*ij*_ is the *j*th voltage step (in mV) in the *i*th pulse in the protocol, and *t*_*i*_ is the time (in ms) following the *i*th pulse in the protocol. We can then optimise the max-med criterion with respect to *ϕ* via CMA-ES to determine an optimal protocol 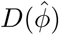. To initialise the CMA-ES optimisation we take 100 random samples from within the voltage and time step bounds defined above ([−50, +40] and [1050, 21000] respectively), and then use the *ϕ* that gives the largest value of the max-med criterion as our initialisation for the optimiser.

### 2.5 Fitting models to synthetic data generated with an optimised protocol

With an optimised protocol we can then generate a new set of synthetic data using the methodology described in section 2.2. Our new protocol is comprised of two multi-step 10 s pulses of interest whereas the Milnes protocol has just one single-step 10 s pulse. We, therefore, generate data for only 5 sweeps (cf. 10) of our new protocol to ensure that the synthetic data under the optimised protocol has the same number of data points as the Milnes synthetic data. In Figure 5 we plot this new synthetic current and proportion open data (right top and bottom respectively) alongside the optimised protocol (left). We can then fit the 15 drug-binding models to this new synthetic data using the same method described in section 2.3, the fits are shown in Figure 6. We see that, when compared to Figure 3, we now have a number of models that do not appear to fit the data very well. This is backed up by Figure 4, where we see a greater spread in maximised log-likelihood values in the optimised protocol case compared to Milnes case, making it easier to distinguish between models. Model 7, the ‘true’ data-generating model, once again has the largest maximised log-likelihood, however, this time no other models have maximised log-likelihoods within 10^5^ of Model 7. In the supplementary material in Figure S2 we include a plot comparing the model parameters fitted to the Milnes data, and the model parameters fitted to the optimised protocol data.

**Figure 5:**
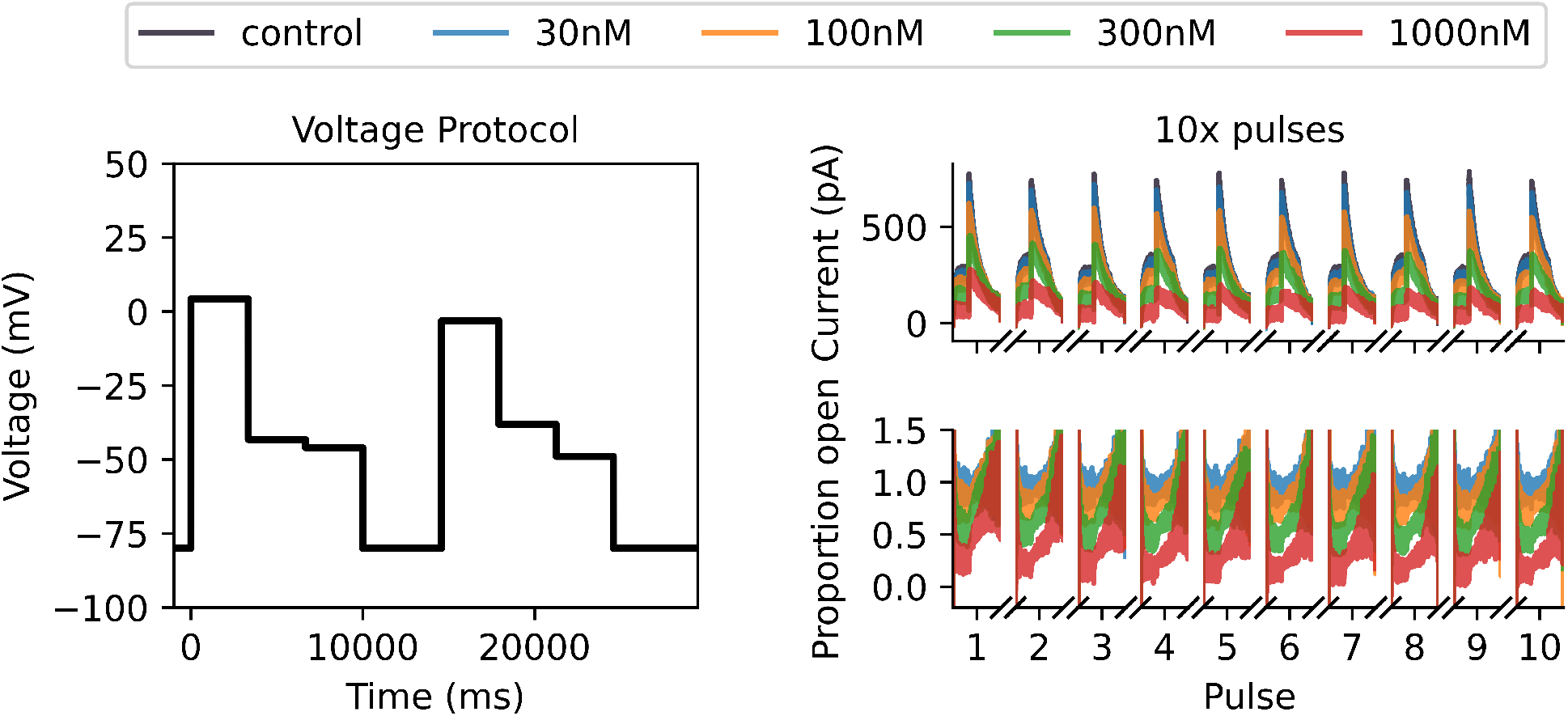
Synthetic data for drug-binding Model 7 generated under an optimised protocol with the Lei 37°C hERG model. At the left we plot our optimised protocol with two 10s pulses each with three optimised voltage steps. At the top right, we plot the control and drug currents for 5 sweeps of the optimised protocol. We only plot the currents that occur during the two 10s pulses per protocol sweep. At the bottom right, we plot the corresponding proportion open for each drug concentration which is calculated by dividing each drug sweep by the control current.

**Figure 6:**
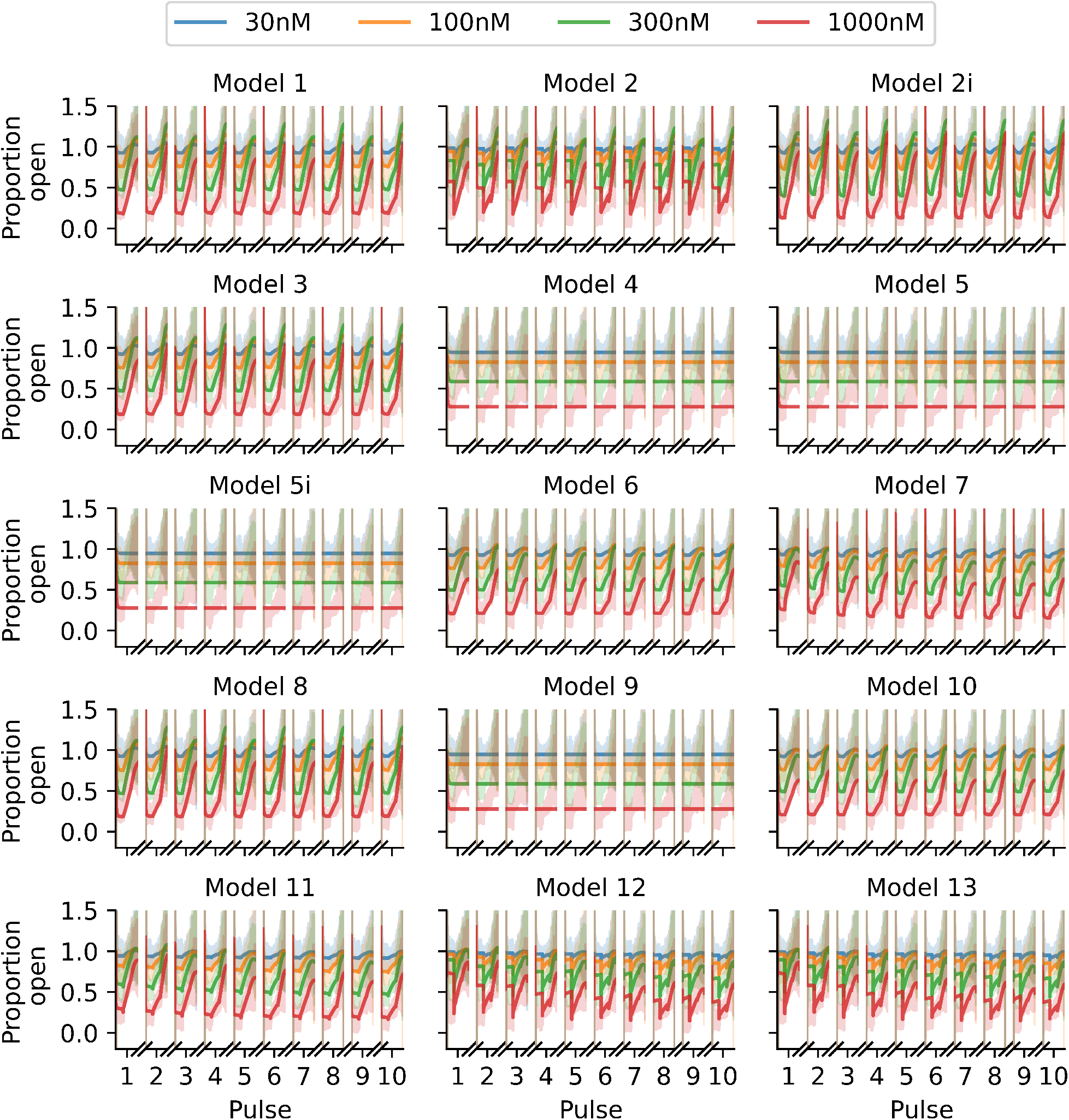
Fits of the binding models to the optimised protocol proportion open synthetic data shown in Figure 5, in the same style as Figure 3. Note how many fits are now visually distinguishable, we can immediately see that many models provide a worse fit to these data than others.

### 2.6 A discrepant hERG model

So far we have been working under the assumption that the structure and parameterisation of the underlying data-generating hERG model is known exactly. Bernardo and Smith [34] describe this as an M-closed model space, where the true data-generating model is included in the set of models considered for model selection. We can never guarantee this practically, so we now introduce an example where the assumed hERG model used in the model fitting and protocol optimisation steps is different from the hERG model used during the data-generating process. We are now operating in an M-open model space where the true data-generating model is not within our set of candidate models.

To generate synthetic data, we now use the Lu model as illustrated in Figure 1b. In Figure 7 we plot a comparison between control currents under the Lu model and the 37°C Lei model previously used to generate synthetic data. We set the conductances (*g*_Kr_) for both models to be equal. We can then repeat the procedure described in sections 2.2, 2.3, 2.4, and 2.5, but this time using the Lu hERG model when generating any synthetic data (Model 7 is still used as the data-generating drug-binding model). In Figure 8 we plot the log-likelihoods obtained using the Lu model as the true data-generating hERG model, but (wrongly) assuming that the 37°C Beattie model is the correct hERG model when fitting the drug-binding models to the data.

**Figure 7:**
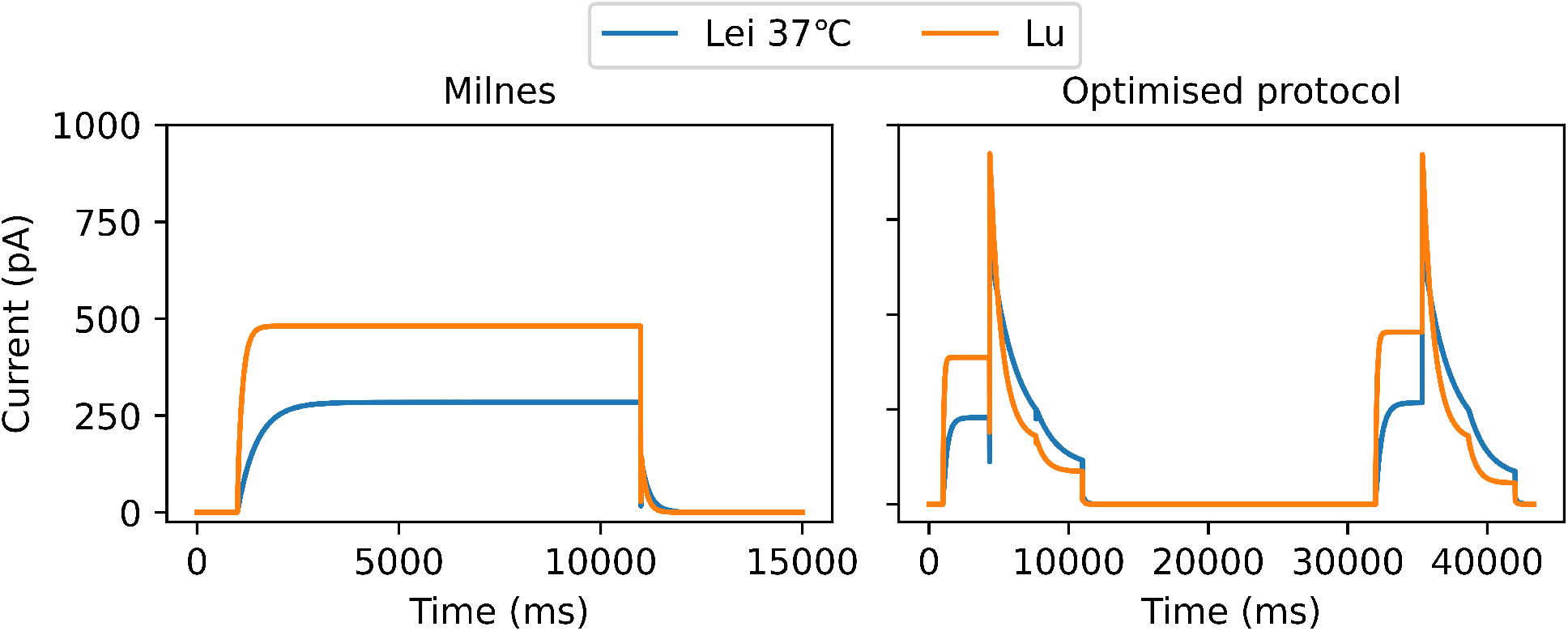
A comparison of control currents between the Lei model (Figure 1a) parameterised for 37°C and the Lu model (Figure 1b). To the left we plot the currents in response to one sweep of the Milnes protocol, and to the right in response to one sweep of the optimised protocol from Figure 5. Conductances have been set to 33.3 *µ*S for both models.

**Figure 8:**
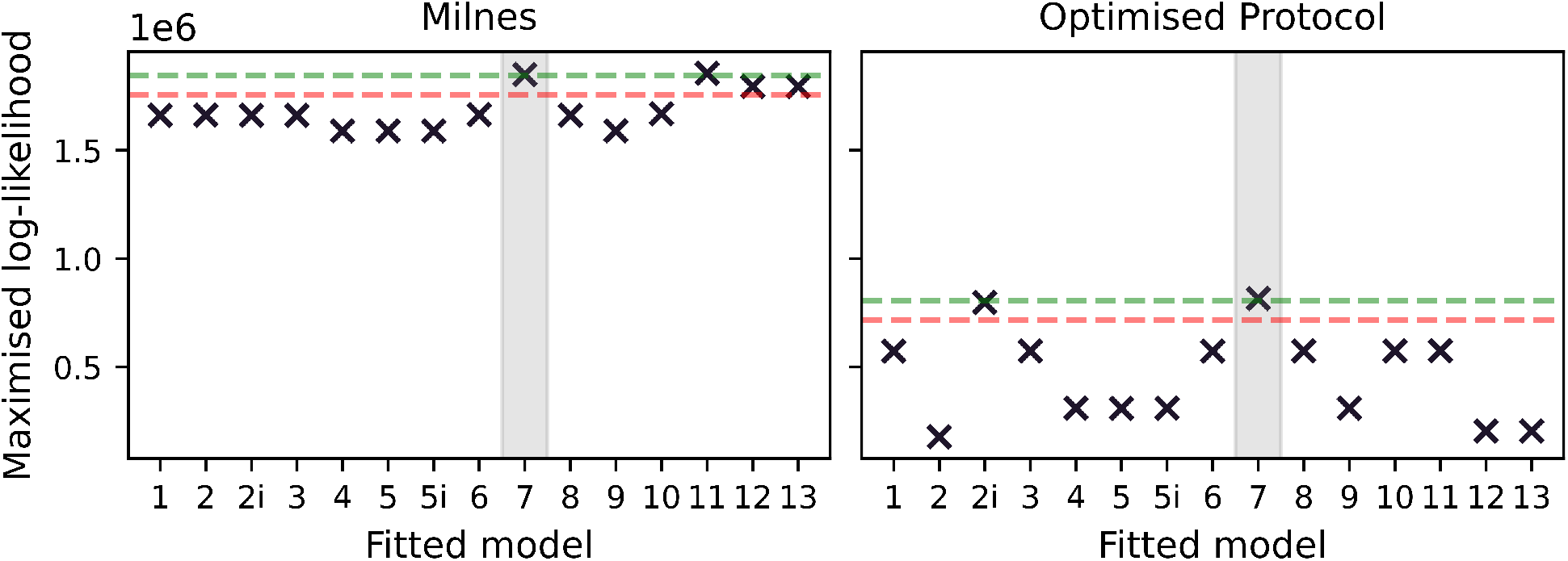
Maximised log-likelihoods for model fits to synthetic data generated under the modified Milnes protocol (left) and an optimised protocol (right). This is for the discrepant hERG model case; the data-generating hERG model is the Lu model, while the Lei 37°C is used for model fitting. In each plot we include a green dashed line at 10^4^ below the largest maximised log-likelihood and a red dashed line at 10^5^ below the largest maximised log-likelihood. We also shade, in grey, around the data-generating binding model.

We appear to get quite similar results to the non-discrepant case, with the optimised protocol once again spreading out the values of the log-likelihoods and pointing towards Model 7 as the data-generating model. However, as we will see in section 3, we find these methods are less effective for other data-generating drug-binding models when discrepancy is introduced. In the supplementary material we include equivalent plots to Figures 2, 3, 5, and 6 in Figures S3, S4, S5, and S6 respectively. In Figure S7, we also include a plot comparing the model parameters fitted to the Milnes data and the model parameters fitted to the optimised protocol data.

## 3 Results

In section 2 we saw how our methodology performed for synthetic data generated under one drug-binding model parameterised for one drug compound. We now repeat these methods across a range of models and drugs to demonstrate how the procedure performs in differing circumstances.

### 3.1 Verapamil: non-trapping dynamics

We begin by considering verapamil again, but this time we generate synthetic data for each of the 15 drug-binding models. Starting with the previous fits to real verapamil data from [18] to generate the synthetic data, in the top half of Figure 9 we plot heatmaps of the log-likelihoods obtained via the section 2 methodology. This is for the case with no hERG model discrepancy. The y-axis represents which drug-binding model is used to generate the synthetic data, and we then get a corresponding log-likelihood for each fitted drug-binding model on the x-axis. We plot both the log-likelihoods obtained for the Milnes protocol data fits (top left), and for the optimised protocol data fits (top right). For each row, we highlight (with green squares) the fitted models within 10^4^ of the largest log-likelihood in that row. We then also highlight (with red squares) the fitted models that are within 10^5^ of the largest log-likelihood (that are not already highlighted in green). Black dots are plotted down the diagonal indicating where the fitted model corresponds to the data-generating model. The first notable observation from this plot is that the number of models within the 10^4^ and 10^5^ thresholds is significantly reduced in the optimised protocol case compared to the Milnes case. We also note that in both cases the full diagonal is within the 10^4^ threshold. Note that there is a large amount of model nesting between drug-binding models, which we illustrate in Figure 10. We consider some model, model A, to be nested within another model, model B, if by fixing one or more of the parameters in model B we can reduce model B to model A. For this reason, we expect that (in this non-discrepant hERG case) if model A is nested within model B, and model A is the true data-generating model, then model B should also be able to fit the data at least equally well (perhaps even slightly better given higher complexity and more parameters), so the maximised log-likelihoods for Model A and B should be roughly equivalent. At the top of the tree diagram are the models that are not nested within any others; models 7, 10, 11 and 13. If we consider the heatmap in the top right of Figure 9 corresponding to the optimised protocol, we see that for synthetic data generated under models 7, 10, and 11, the only fitted models with a log-likelihood within the 10^4^ threshold is the true data-generating model. Model 13, the other model with no nesting, only has one other model within this 10^4^ threshold (model 12). Considering the nested models in Figure 10, we can explain some of the cases where multiple models sit within the 10^4^ threshold. In the optimised protocol case in the top right of Figure 9, the 10^4^ threshold highlighted squares in the heatmap rows corresponding to models 2, 2i, 8, and 12 can all be explained by model nesting. On the other hand, data-generated under models 1, 3, 4, 5, 5i, 6, and 9 all have at least one model within the 10^4^ threshold that cannot be explained by nesting.

**Figure 9:**
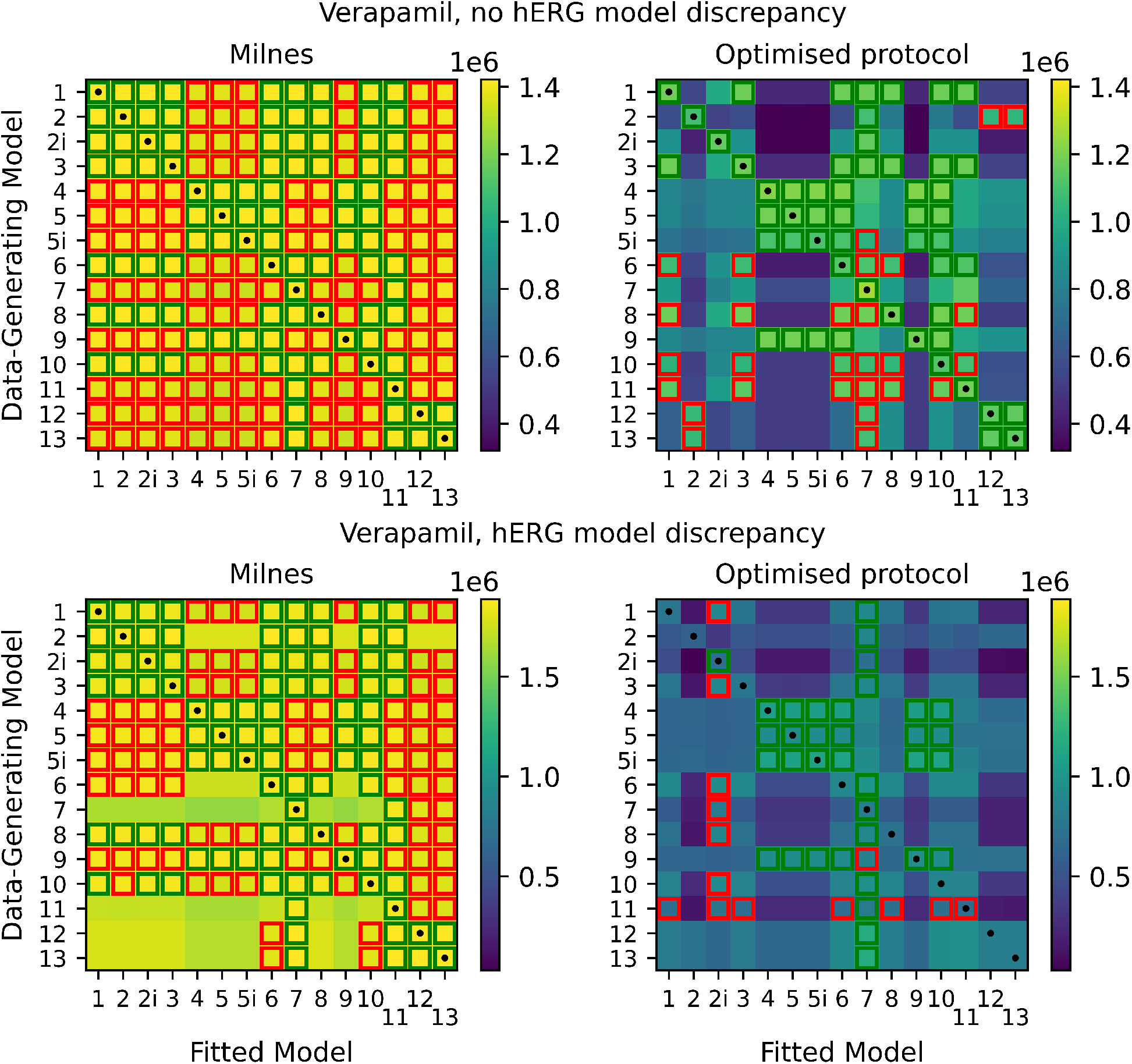
Top: heatmaps of maximised log-likelihoods for fitted models to Milnes and optimised protocol synthetic verapamil data. The fitted models within 10^4^ of the largest log-likelihood (very good fits) in each row are highlighted in green squares. The fitted models that are between 10^4^ and 10^5^ below the largest log-likelihood (reasonable fits) in each row are highlighted in red squares. Black dots are plotted down the diagonal of each heatmap where the fitted model corresponds to the data-generating model. Note that our optimal protocol results in a far bigger spread of maximised log-likelihoods. Bottom: equivalent heatmaps of maximised log-likelihoods with discrepancy between the data-generating hERG model and the hERG model used to fit the data. The introduced hERG model discrepancy makes identifying the ‘true’ data-generating drug-binding model difficult.

**Figure 10:**
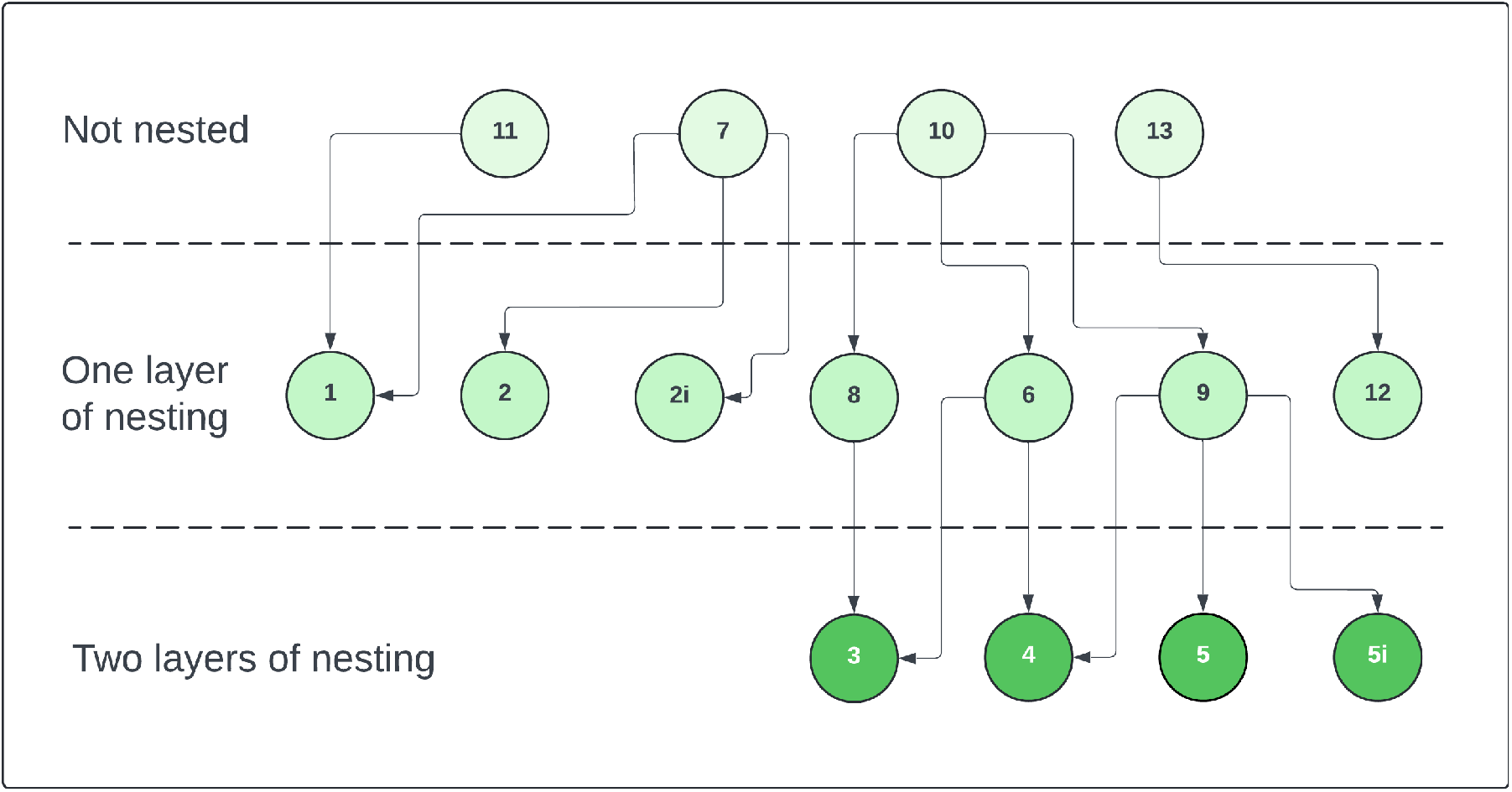
Graph illustrating nesting between drug binding models. An arrow pointing from model A to model B indicates that model B is nested within model A.

We now switch to the discrepant hERG model case where synthetic data were generated using the Lu physiological hERG model but we use the Lei model in fitting. In the bottom half of Figure 9, we plot equivalent heatmaps to those at the top of the plot, but for the discrepant case. The purpose of considering this case, is to address the more realistic scenario where we do not have a perfect model of hERG channel dynamics. We notice that the number of highlighted 10^4^ and 10^5^ threshold squares is once again significantly reduced in the optimised protocol case. However, in most cases, the best-fitting binding model is no longer the one that generated the data. As a result, the diagonal is highlighted significantly less than in the non-discrepant case, with only 6 models (2i, 4, 5, 5i, 7, and 9) sitting within the 10^4^ threshold. Clearly, the hERG model discrepancy is causing issues with identifying the true binding mechanisms.

### 3.2 Bepridil: trapping dynamics

We can repeat the process described in the previous section for a different drug, this time with observed trapping behaviour [10, 11]. In the supplementary material we include, in Figure S8, an example of model output for a drug with trapping behaviour, compared to one with no trapping behaviour. In the top half of Figure 11, we plot the heatmaps of maximised log-likelihoods for the non-discrepant case as previously. The optimised protocol heatmap is very similar to that obtained with verapamil; the highlighted squares in the rows corresponding to models 1, 2, 2i, 3, 5, 5i, 8, and 9 are identical, while there are only a couple of differences for each of the other models. We can also again consider the discrepant hERG case, and we plot the heatmaps for this in the bottom half of Figure 11. Again this shows relatively similar results to those seen with verapamil; when we have discrepancy in the hERG model, it becomes difficult to determine the true data-generating drug-binding model. In the supplementary material, we also include heatmaps in Figure S9 for results obtained for chlorpromazine, a fast-binding drug with a suspected open-binding preference (over inactive-binding). These results appear similar to those obtained for verapamil and bepridil.

**Figure 11:**
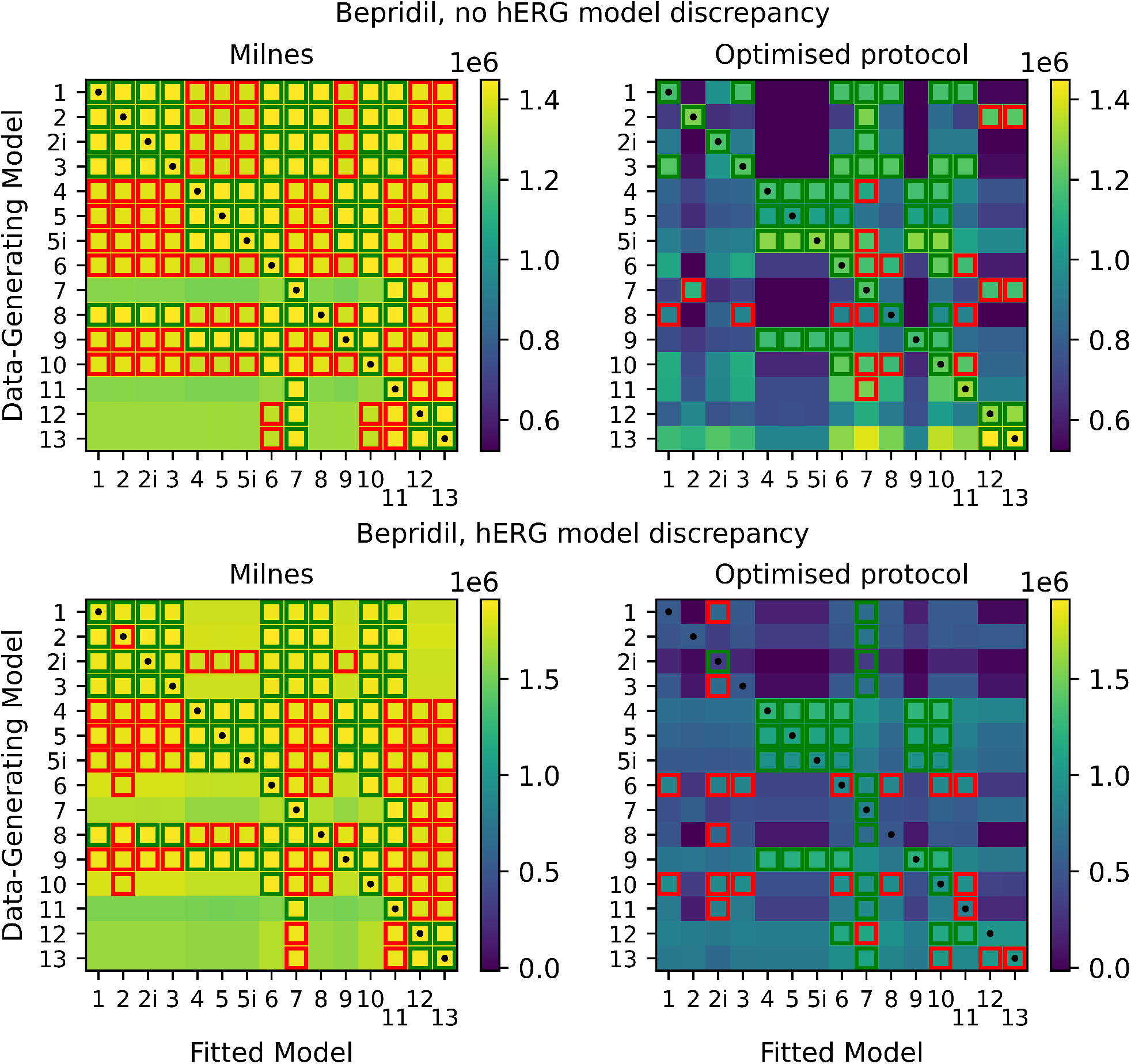
Top: heatmaps of maximised log-likelihoods for fitted models to Milnes and optimised protocol synthetic bepridil data. Bottom: equivalent heatmaps of maximised log-likelihoods with discrepancy between the data-generating hERG model and the hERG model used to fit the data.

### 3.3 Dofetilide: slow-binding dynamics

Bepridil and verapamil both have relatively fast binding dynamics, so we also consider the slow-binding drug dofetilide [35]. In the supplementary material, in Figure S8, we include an example of model output for a drug with slow dynamics, compared to one with fast dynamics. Once again, we plot the heatmaps of maximised log-likelihoods as shown in Figure 12, and we see that nearly all models can fit the Milnes data well. Unlike the fast dynamic drugs, with dofetilide, we get much less of a reduction in 10^4^ and 10^5^ threshold squares with the optimised protocol (in both the non-discrepant and discrepant cases). It is not obvious what causes this difference between fast and slow-binding drugs and it may indicate that our median optimisation objective function is less effective in the slow-binding case.

**Figure 12:**
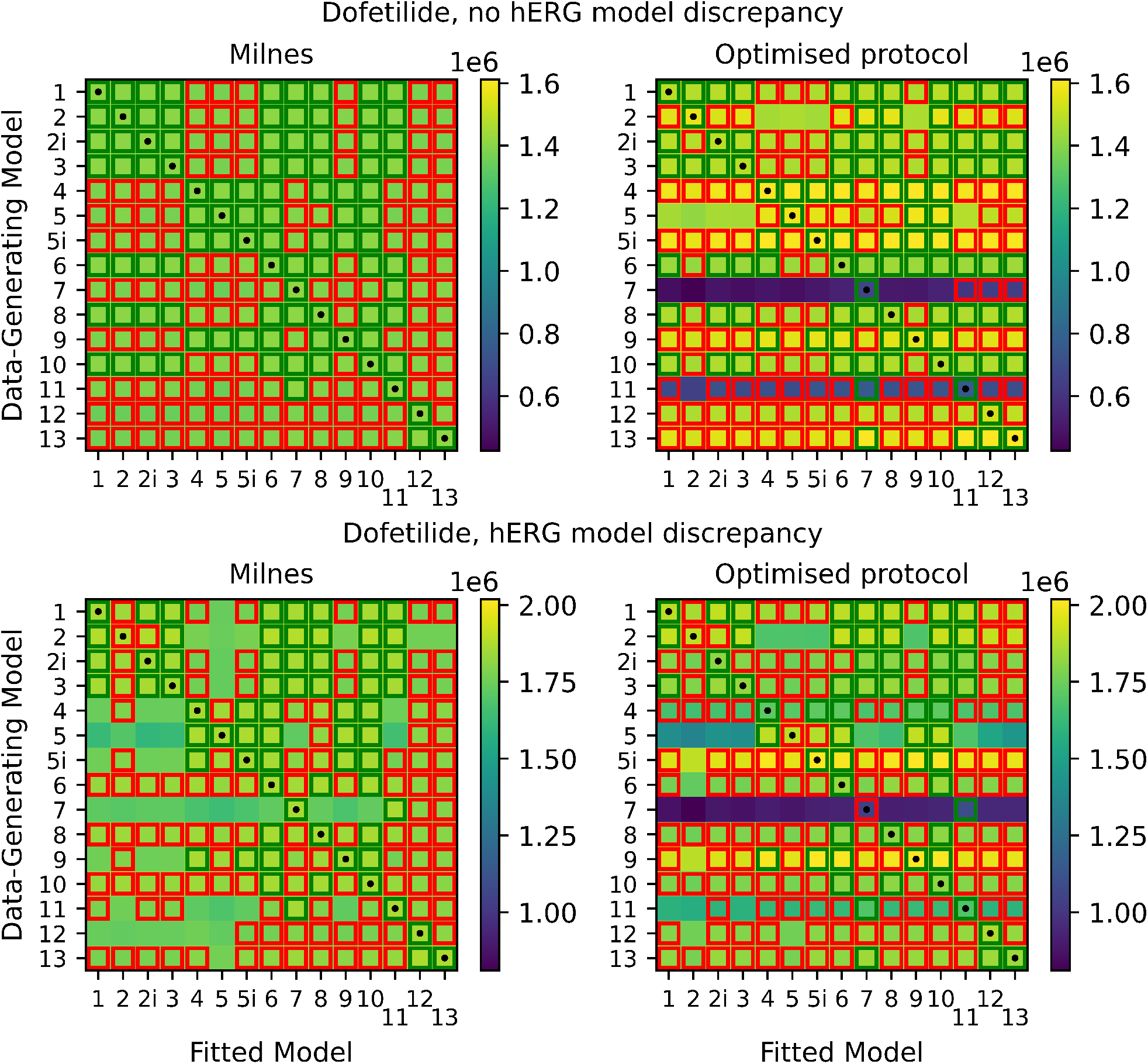
Top: heatmaps of maximised log-likelihoods for fitted models to Milnes and optimised protocol synthetic dofetilide data. Bottom: equivalent heatmaps of maximised log-likelihoods with discrepancy between the data-generating hERG model and the hERG model used to fit the data.

## 4 Discussion

In this study, we have outlined a methodology for generating optimised voltage protocol designs to assist in distinguishing between models of drug-binding dynamics. By undertaking a synthetic data study, we have seen the potential benefits of this methodology when the true physiological model of hERG is known. Log-likelihoods of models fitted to data generated under our optimised protocols indicate a divergence in the quality of fits when compared to fits to data generated under a simple Milnes protocol. This suggests that this OED procedure could assist in establishing the true binding dynamics of a compound. The method was less effective when considering synthetic data emulating a slow-binding drug, dofetilide, when compared to drugs with fast dynamics such as verapamil and bepridil. We also considered how discrepancy in the hERG model used to fit the data (compared to the data-generating hERG model) influenced the outcomes of our methodology. We found that when we used the Lu hERG model to generate synthetic data and the Lei 37°C hERG model to fit models to this data, log-likelihoods of model fits indicated that establishing the true data-generating drug-binding model became more difficult. This suggests that the underlying hERG model does play an important role when fitting drug-binding models to data and stresses the importance of continuing to improve basic models of physiological channel behaviour. The proposed methodology could perhaps be improved by considering fitting a hERG model to the obtained control currents before fitting the drug-binding models (or fitting both the hERG and drug-binding models simultaneously).

We note that our results depend on the initial drug-binding model parameterisations for each of the three drug compounds, which come from model fits obtained by Lei *et al*. [18]. The quality of these model fits was quite variable from model to model and from compound to compound. This represents a limitation of a synthetic data study and motivates trialling the methodology on real data.

The methods used to fit the drug-binding models in this paper were developed to improve on those used by Lei *et al*. and others [18, 26, 12]. While we have similarly fitted our models to the proportion of hERG in the open state, the log-likelihoods derived from the normal ratio PDF described in section 2.3 differ from the simple weighted sum of squares method used previously. This likelihood fitting method accounts for the true noise model in the data-generating process (in this synthetic scenario), and it also allows us to fit the binding models to the full data sweep. In Lei *et al*. [18], the weighted sum of squares method required low currents at the start of each 10s pulse to be cropped out to prevent noisy open proportion data from biasing the fit.

After some consideration and testing, the max-med optimality criterion was chosen over other possible alternatives such as maximising the mean or the minimum of the pairwise sum of squares difference between model current traces. Using the mean or minimum, as opposed to the median, tended to result in one or two model pairwise differences biasing the objective function score and leaving many of the other current traces indistinguishable from each other. It would be useful to perform more rigorous comparisons between optimality criteria to see if there are scenarios where alternatives are more effective.

In section 3.1, we noted the presence of nesting between the 15 binding models. This nesting suggests that perhaps reducing our optimisation problem to consider only pairwise differences between the non-nested models (7, 10, 11, and 13) could be an alternative starting point given all other nested models are simplified versions of these.

To conclude, the proposed OED methodology shows promise in determining the true binding dynamics, but care must be taken to ensure that we have a well-calibrated model of hERG current if applied to a real data study.

## Data Availability

Open source code is freely available at: www.github.com/CardiacModelling/binding_model_OED.

## Acknowledgements

Model fitting and experimental design optimisations were performed using the University of Nottingham’s on-premise High Performance Computing (HPC) service.

