## Supplementary Material for "Optimising experimental designs for model selection of ion channel drug binding mechanisms"

### Supplementary Figures

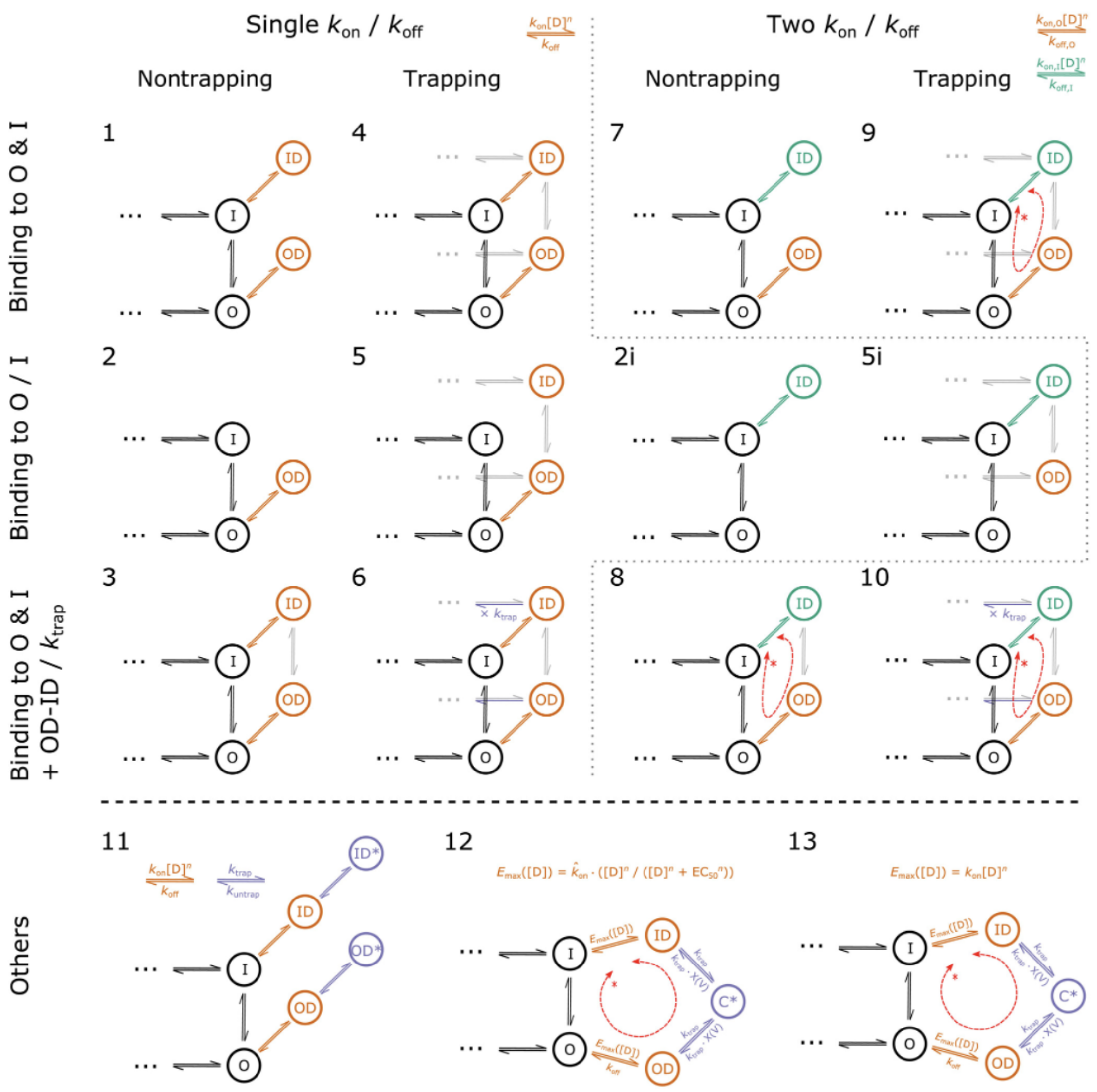


Figure S1: Markov diagrams of pharmacological drug-binding models from Lei et al. [1] reproduced here under a CC-BY licence. Trapping components are indicated as grey 'mirror images' of the hERG physiological model.

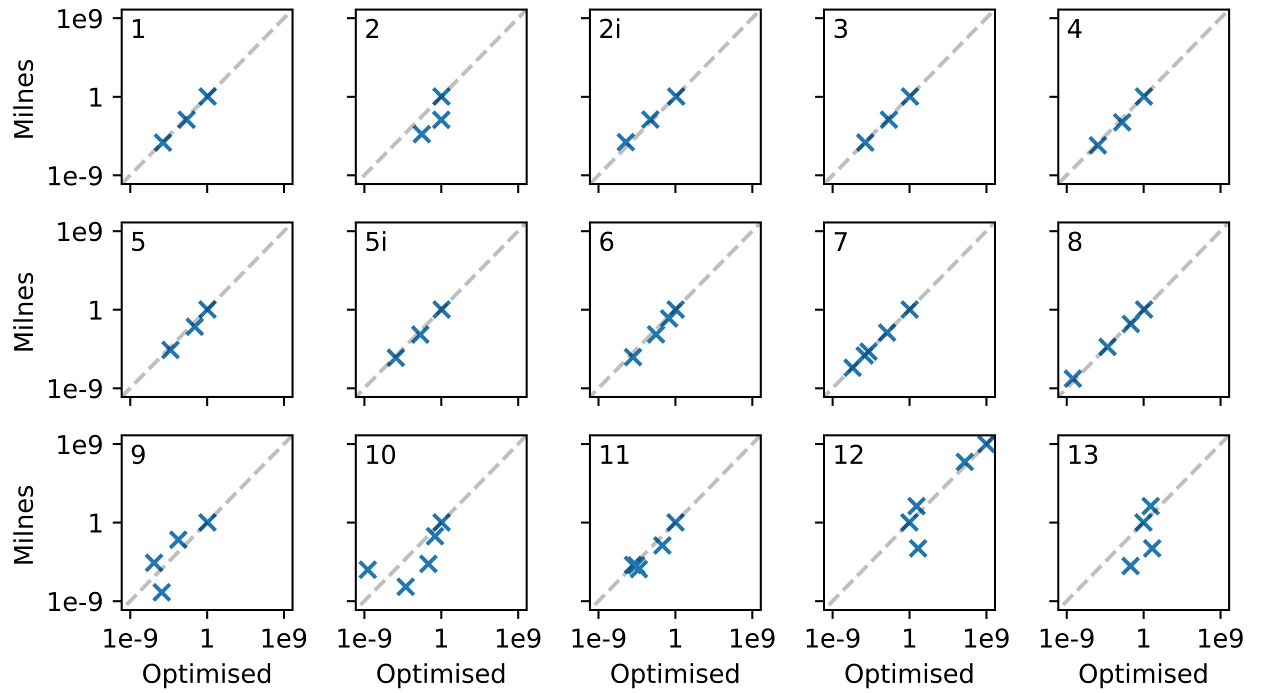


Figure S2: Comparison plots of model parameters fitted to Milnes data and optimised protocol data. This is for the verapamil model 7 case, with no model discrepancy. The blue crosses indicate the values of the fitted model parameters for each binding model.

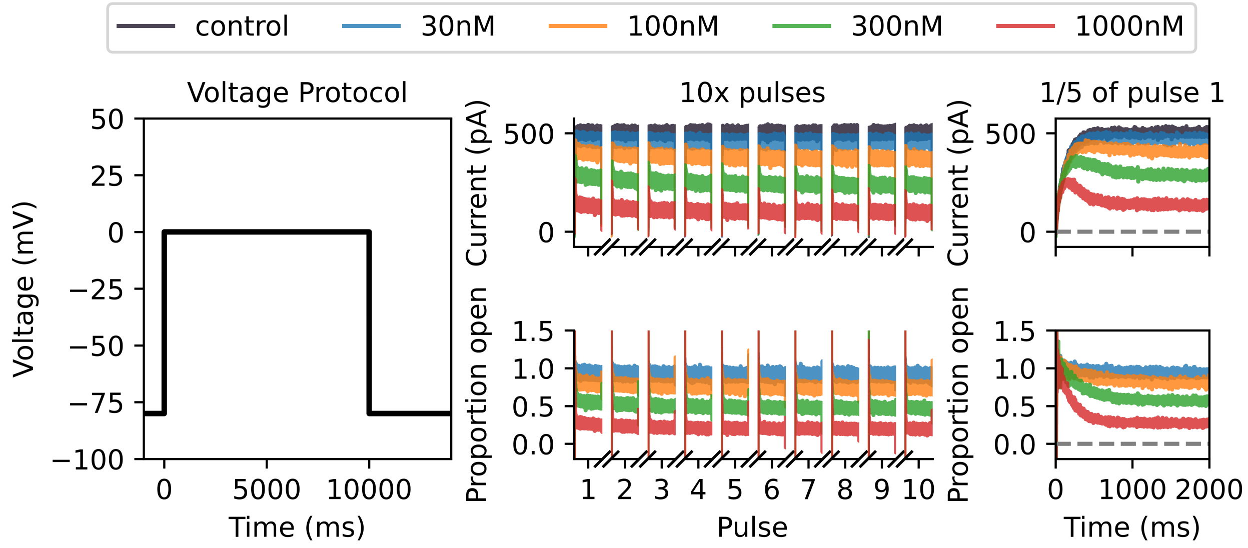


Figure S3: Synthetic verapamil data under drug-binding model 7 generated under a Milnes protocol with the Lu hERG model. At the left, we plot a single sweep of the Milnes protocol. In the top middle, we plot the control and drug currents for 10 sweeps of the Milnes protocol. We only plot the currents that occur during the 10s pulse at 0 mV for each sweep. In the bottom middle, we plot the corresponding proportion open for each drug concentration which is calculated by dividing each drug sweep by the control current. On the right, we include a zoomed-in look at the first $\frac{1}{5}$ (2 s) of the first pulse for both currents and proportion open.

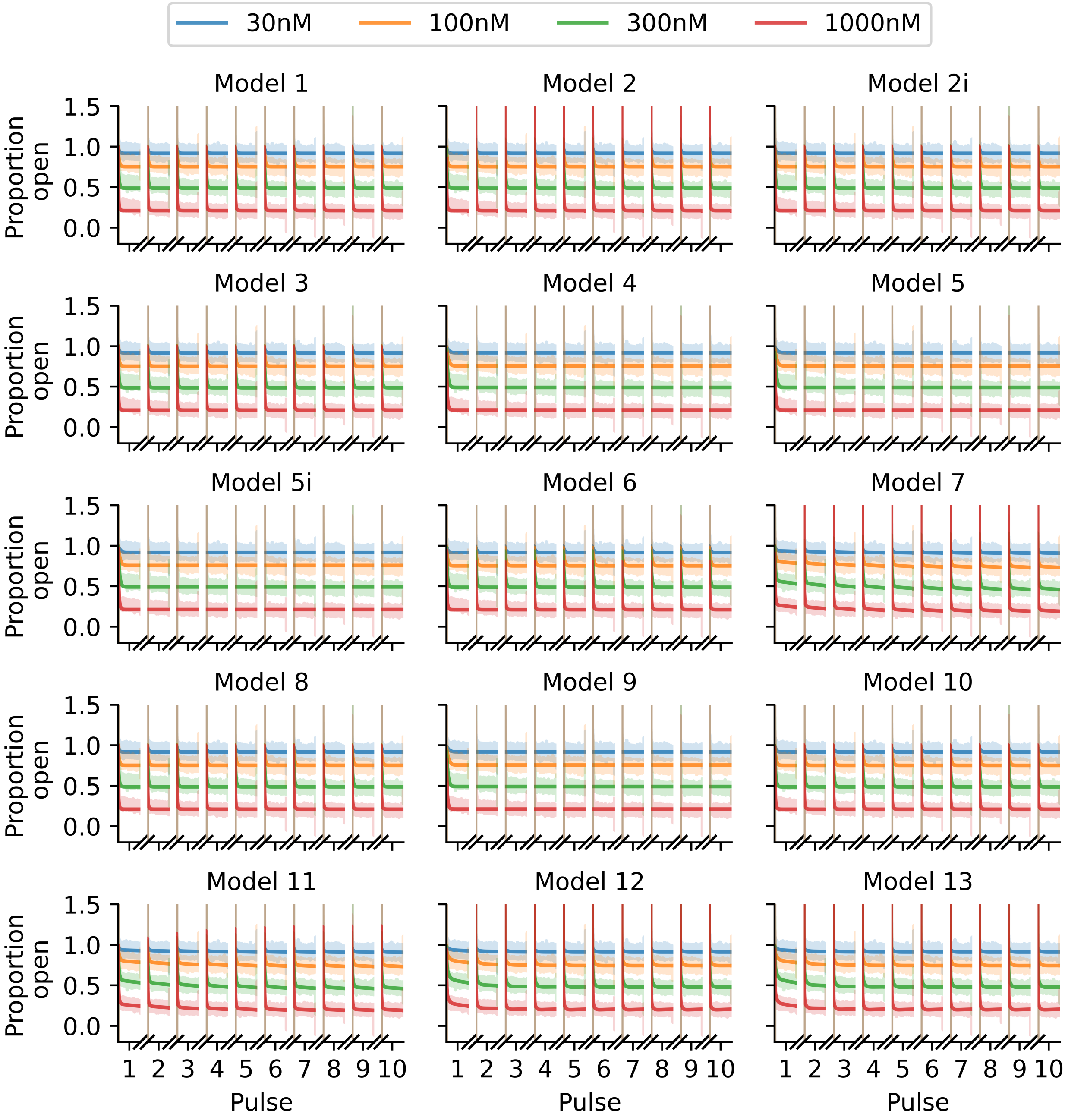


Figure S4: Fits of the 15 binding models to the Milnes protocol proportion-open synthetic data shown in Figure S3. The fitted models are shown with solid lines, and we plot the data slightly shaded out behind these fits. While the data is generated using the Lu hERG model, fits are obtained assuming the Lei 37°C hERG model.

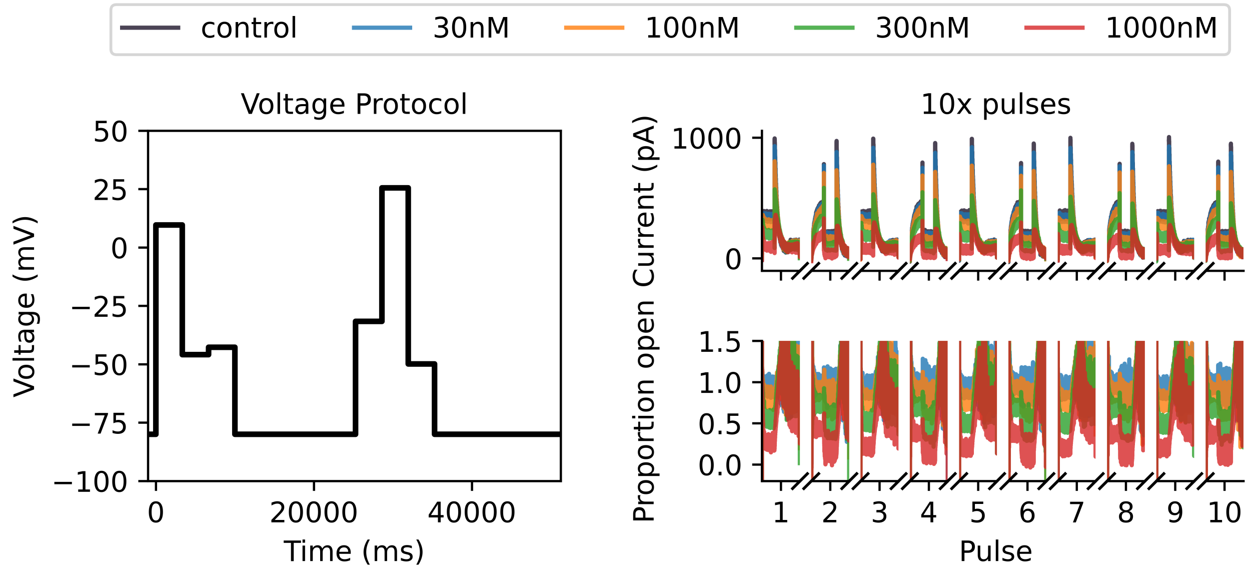


Figure S5: Synthetic data for drug-binding model 7 generated under an optimised protocol with the Lu hERG model. At the left we plot our optimised protocol and with two 10s pulses each with three optimised voltage steps. At the top right, we plot the control and drug currents for 5 sweeps of the optimised protocol. We only plot the currents that occur during the two 10s pulses per protocol sweep. At the bottom right, we plot the corresponding proportion open for each drug concentration which is calculated by dividing each drug sweep by the control current.

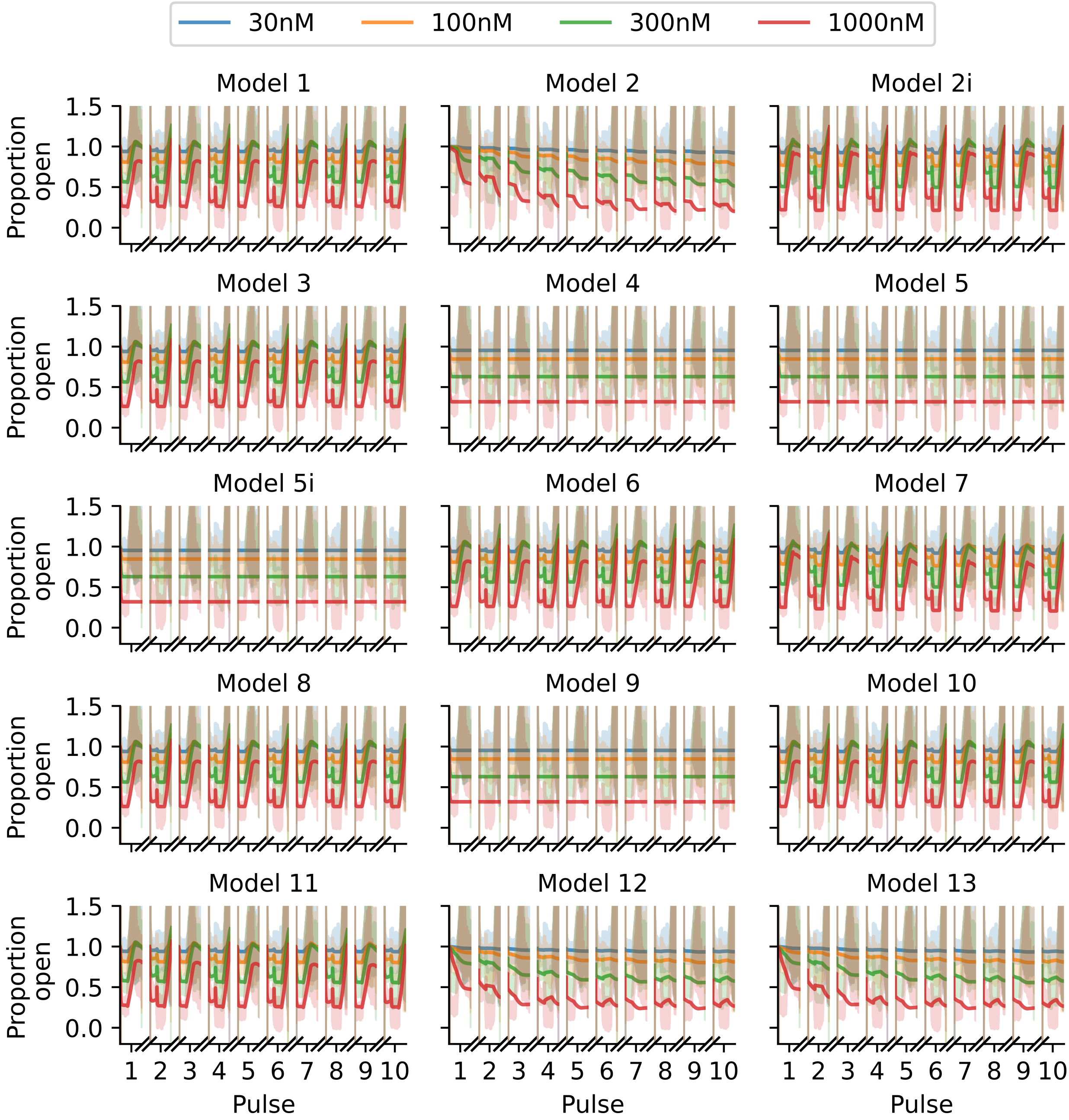


Figure S6: Fits of the binding models to the optimised protocol proportion open synthetic data shown in Figure S5, in the same style as Figure S4.

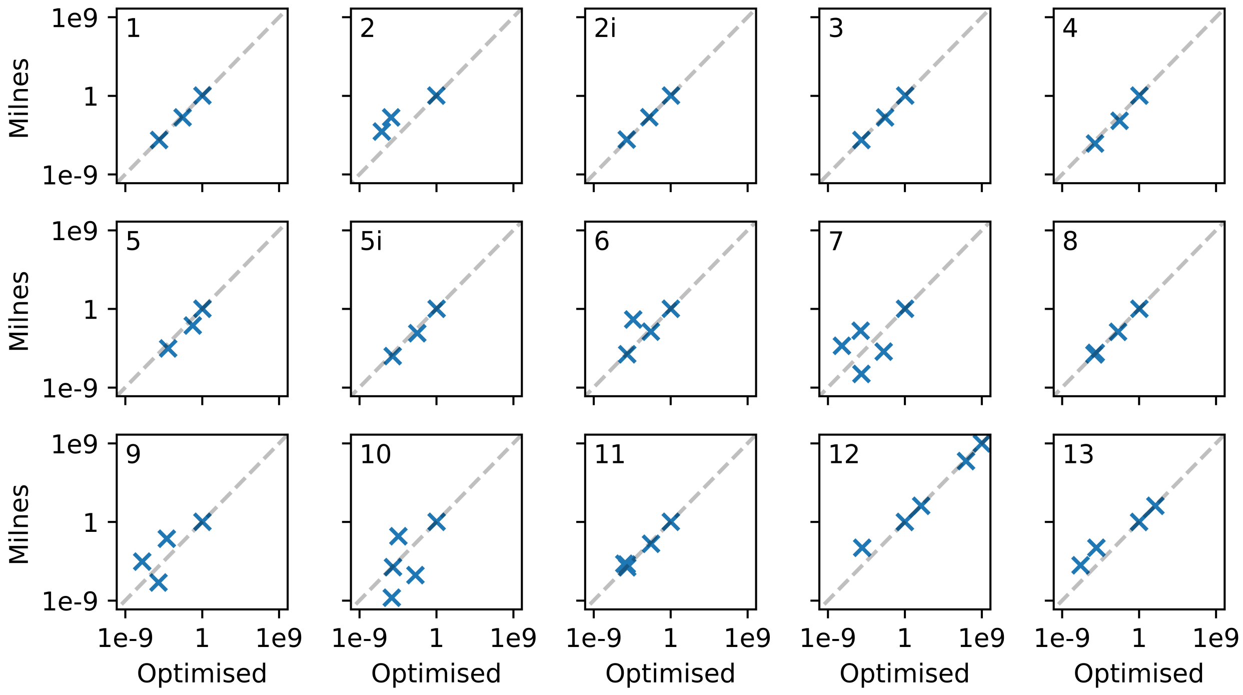


Figure S7: Comparison plots of model parameters fitted to Milnes data and optimised protocol data. This is for the verapamil model 7 case, with discrepancy between the data-generating hERG model and the hERG model used for model fitting. The blue crosses indicate the values of the fitted model parameters for each binding model.

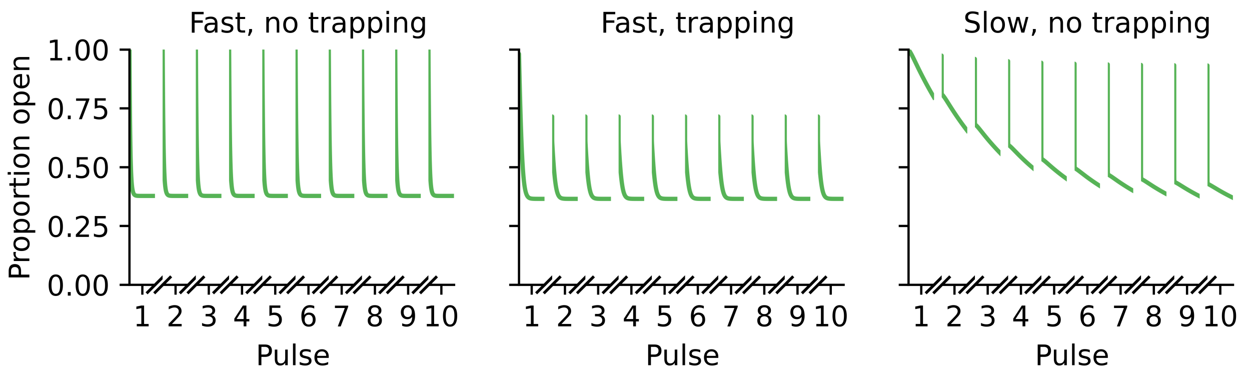


Figure S8: Model output for example drugs with different binding dynamics under 10 sweeps of the Milnes protocol. The left plot illustrates fast binding dynamics with no trapping behaviour; the drug quickly reduces the proportion open to 0.4 on each pulse but returns to being completely open between pulses. The middle plot also shows fast binding, but with trapping behaviour; this time the channel does not completely open between pulses. In the right plot, we illustrate slow binding with trapping behaviour; about 7 pulses of the Milnes protocol are required for the open proportion to reach an equilibrium at 0.4.

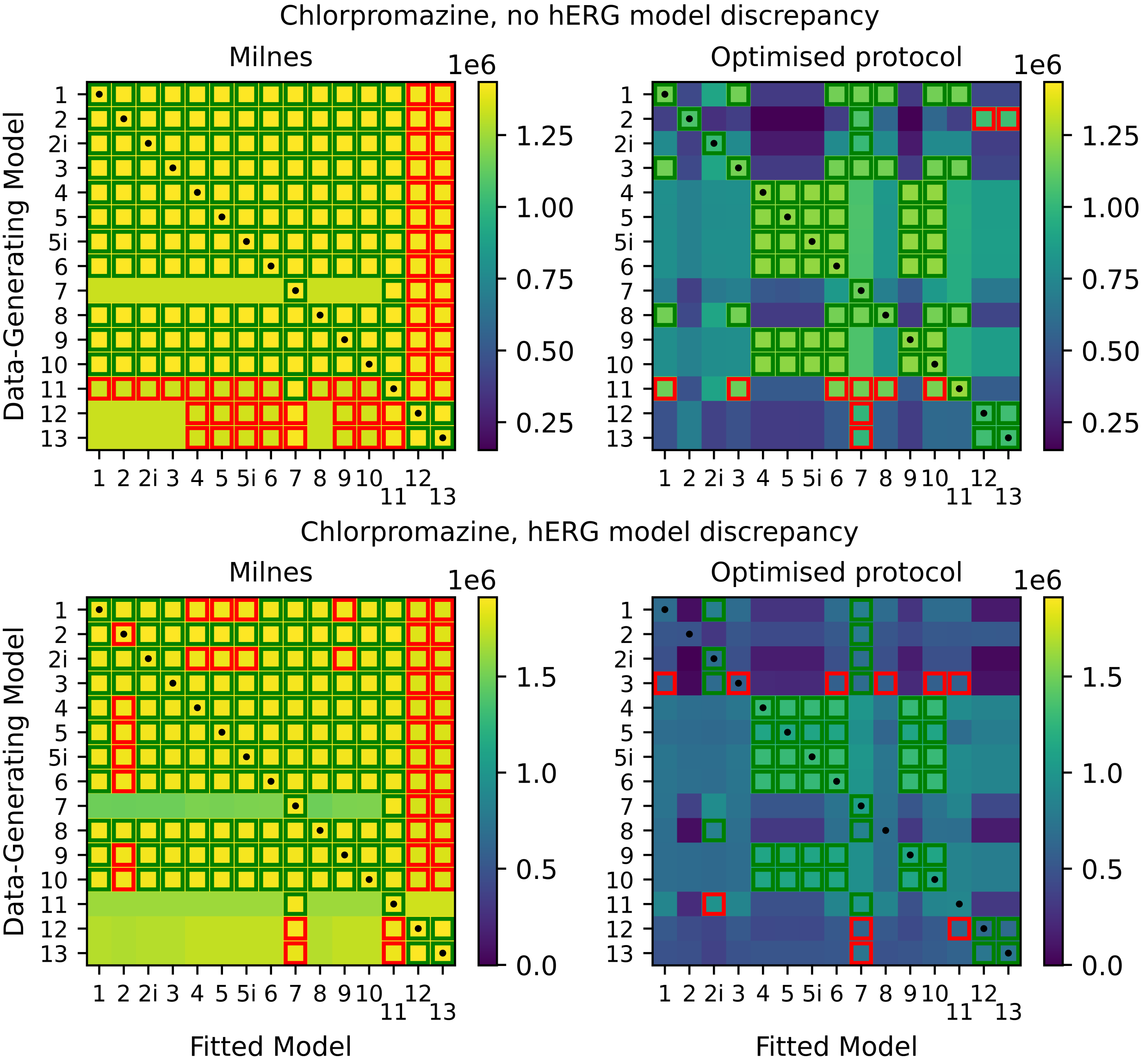


Figure S9: Top: heatmaps of maximised log-likelihoods for fitted models to Milnes and optimised protocol synthetic chlorpromazine data. Bottom: equivalent heatmaps of maximised log-likelihoods with discrepancy between the data-generating hERG model and the hERG model used to fit the data.
